# No two are the same: Identification and characterization of novel hyperactive PrfA* variants in *Listeria monocytogenes*

**DOI:** 10.64898/2026.10.13.758832

**Authors:** Maja Andiel, Caroline Borreby, Lea Füllgrabe, Jasmin Petrovac, Birgitte H. Kallipolitis, Jeanine Rismondo

## Abstract

The virulence potential of the human pathogen *Listeria monocytogenes* is primarily controlled by the master virulence regulator PrfA. PrfA activity is tightly regulated to prevent virulence gene expression outside the host, while enabling their rapid induction upon host cell entry. One mechanism controlling PrfA activity is the sugar-dependent inhibition. Readily metabolizable sugars, such as glucose or cellobiose, inhibit PrfA; however, the precise mechanism and the factors involved in this regulation remain unknown. Here, a suppressor screen was performed to identify these factors. Although the screen did not reveal any factors mediating the sugar-dependent inhibition of PrfA, it resulted in the isolation of 13 novel hyperactive PrfA variants, termed PrfA*. All newly identified PrfA* variants were characterized using growth, PlcB activity and promoter activity assays. Interestingly, two of these PrfA* variants, PrfA^Y126F^ and PrfA^V153M^, were shown to function even in the absence of PrfA’s cofactor glutathione. In addition, a subset of variants was analyzed for intracellular replication using *in vitro* infection assays. Across all assays, the effect of the PrfA* mutations were found to be highly variant- and gene-dependent, with no two variants exhibiting identical phenotypes. Importantly, these results demonstrate that the transcriptional and phenotypic consequences of PrfA* mutations cannot be reliably predicted from a single assay. Thus, comprehensive characterization using multiple complementary approaches is essential to determine the effects of PrfA* mutations on the expression of PrfA-regulated genes and the pathogenic potential of *L. monocytogenes* strains carrying these mutations.

**Author summary:** The activity of the main virulence regulator PrfA of the human pathogen *L. monocytogenes* is regulated at multiple levels. One intriguing mechanism is the sugar-dependent inhibition of PrfA, which was identified more than three decades ago but still remains poorly understood. We used a suppressor screen to identify factors involved in this regulation. Surprisingly, the screen identified 13 novel hyperactive PrfA variants that were less inhibited by cellobiose. In-depth characterization revealed that although all variants increased PrfA activity, each variant produced distinct phenotypes, with effects that depended on both the specific amino acid substitution and the PrfA-regulated gene. Our findings further indicate that individual PrfA mutations can alter functional properties of the regulator, including its structural state and its cofactor dependence. Remarkably, enhanced PrfA activity has now been associated with mutations at 24 of its 237 amino acid positions. This broad spectrum of activating mutations may explain why genetic screens aimed at identifying factors involved in sugar-dependent PrfA inhibition repeatedly yielded hyperactive PrfA variants rather than the factors underlying this regulation.

## Introduction

*Listeria monocytogenes* is a Gram-positive saprophyte ubiquitously found in nature that is able to transition into a human intracellular pathogen upon ingestion of contaminated food products such as raw milk, meat, fish, ready-to-eat products as well as fruits and vegetables [1,2]. Within the human host, *L. monocytogenes* crosses the intestinal epithelium and can cause listeriosis, which in otherwise healthy individuals most commonly results in a mild, self-limiting gastroenteritis. However, in at-risk groups, such as immunocompromised individuals and pregnant women, listeriosis can manifest as severe systemic disease and neonatal infections, respectively [2]. Despite appropriate antibiotic treatment, listeriosis entails a high mortality rate of approximately 20-30% [2,3].

The transition from a saprophytic to a pathogenic lifestyle as well as the overall virulence of *L. monocytogenes* is controlled by the master virulence regulator PrfA (positive regulatory factor A) [4,5]. PrfA activates the expression of virulence genes upon entry into the host. The core virulence regulon comprises ten genes, *plcA, plcB, hly, mpl, actA, orfX, inlA, inlB, inlC*, and *hpt*, which are required for host cell invasion, intracellular growth and cell-to-cell spread [6]. Host cell invasion is mediated by the two internalins InlA and InlB, which interact with the host receptors E-cadherin and Met, respectively, to promote receptor-mediated uptake [7,8]. After endocytosis, *L. monocytogenes* is engulfed in a phagocytic vacuole, which is subsequently lysed by the pore-forming toxin listeriolysin O, as well as a phosphatidylinositol-specific phospholipase C and a broad-range phospholipase C, which are encoded by *hly*, *plcA* and *plcB*, respectively [9,10]. *L. monocytogenes* usually metabolizes glycerol and hexose phosphates from the host cytosol for rapid intracellular growth, latter of which are taken up via the hexose phosphate transporter Hpt [11,12]. Intracellular movement of *L. monocytogenes* as well as cell-to-cell spread is enabled by the actin assembly-inducing protein ActA, which induces the polymerization of host actin molecules into filaments forming an actin tail [13].

The constitutive expression of virulence factors is an energetic burden for *L. monocytogenes*, leading to reduced fitness. Thus, the activity of PrfA needs to be tightly regulated [5,14,15]. On a post-transcriptional level, PrfA production is controlled by an RNA thermosensor found in the 5’-untranslated region of the *prfA* mRNA, which only allows translation at temperatures ≥ 37°C signaling successful host cell invasion [16]. On a post-translational level, PrfA requires a small cofactor to enable DNA binding as it structurally belongs to the family of cAMP receptor protein (Crp) transcription factors [17]. Glutathione has been identified as the cofactor of PrfA, which can be either host-derived or self-synthesized [18–20]. The binding of glutathione results in a conformational change leading to an optimal fold of the helix-turn-helix (HTH) motif enabling DNA binding [18,20]. Furthermore, PrfA activity can be inhibited by the presence of readily metabolizable sugars that are imported via phosphoenolpyruvate (PEP)-dependent transport systems (PTS) such as glucose, fructose and mannose. This inhibitory effect is especially pronounced when *L. monocytogenes* is grown in the presence of the plant-derived sugar cellobiose [21–23]. PTS-independent carbon sources such as glucose-1-phosphate or glycerol do not exert this effect [12,24]. Although this phenomenon has been studied for over 30 years, the precise mechanism of sugar-dependent inhibition of PrfA is still not fully understood. In an attempt to identify factors involved in this mechanism, a suppressor screen was performed. For this, *L. monocytogenes* strains harboring a chloramphenicol resistance cassette under the control of either the promoter of *actA* or *hly* were constructed. These strains were grown on LB plates containing cellobiose and chloramphenicol, a condition in which PrfA is usually inhibited. Therefore, the strains were only able to survive if they acquired mutations that lead to activation of PrfA. While no factors involved in sugar-dependent inhibition were found, 13 novel hyperactive PrfA variants, also referred to as PrfA*, were identified. Here, we characterized the phenotypic consequences of these PrfA* variants. First, the growth of strains harboring the wild type or mutated PrfA variants was compared in LSM using either glucose or glucose-1-phosphate as sole carbon source, latter of which depends on the PrfA-dependent expression of *hpt* [11]. Furthermore, the impact of the PrfA mutations on PrfA activity at low temperatures and in the presence of cellobiose was assessed. Additionally, promoter studies were used to investigate the glutathione dependency of the PrfA* variants. Lastly, intracellular growth of a selection of PrfA* variants was examined. Surprisingly, each PrfA* variant exhibited a distinct phenotype and no two strains behaved identically.

## Results

### 13 novel PrfA* variants were identified in suppressor screens

As a strategy to identify factors involved in sugar-dependent inhibition of the major virulence regulator PrfA, a suppressor screen was performed. For this purpose, a chloramphenicol resistance gene was put under the control of the *actA* and *hly* promoter and integrated into the genome of the *L. monocytogenes* wild type strain EGD-e yielding strains LJR332 and LJR333, respectively. The expression of both genes is controlled by PrfA and thus, LJR332 and LJR333 are only resistant to chloramphenicol when PrfA is active. Plating of high-density cultures of both strains on LB agar containing activated charcoal, egg yolk and chloramphenicol results in bacterial lawn formation; however, only single colonies appeared when the agar additionally contained 25 mM cellobiose. For subsequent analysis, single colonies displaying a whitish halo were selected, as this phenotype indicates production of the lecithin-cleaving enzyme PlcB [25]. The expression of *plcB* is regulated by PrfA and thus, halo formation indicates that PrfA is active. Previous studies already reported the isolation of hyperactivating mutations in PrfA [26–32]. Therefore, the *prfA* gene of 64 halo-producing suppressors was analyzed via Sanger sequencing. In total, 36 suppressors had amino acid substitutions at positions, which were already shown to result in hyperactivity of PrfA, such as I45S, A94T, L140F, G145S, G145C, G155S and P219S (Table S3) [26–32]. Interestingly, 20 strains acquired mutations in *prfA*, which have not been characterized so far (Table S3). Additionally, three strains were isolated in growth assays, which showed improved growth in LSM glucose-1-phosphate or glucose-6-phosphate. Sanger sequencing revealed a substitution of phenylalanine with leucine at position 29 for two of the strains, and of glutamine with lysine at position 35 for the third strain (Table S3). The impact of a Q35 mutation on PrfA activity has not yet been studied. In contrast, a strain carrying a F29M substitution has been previously isolated; however, whether this mutation affects the sugar-dependent inhibition of PrfA had not been investigated [28]. Interestingly, eight halo-producing suppressors did not acquire mutations in *prfA*. Whole genome sequencing analysis was conducted for five of these strains and revealed point mutations in the RNA polymerase subunit encoding genes *rpoB* and *rpoC*, specifically *rpoB^S1128F^*, *rpoB^1086E^, rpoC^R328H^, rpoC^V1060G^* and *rpoC^H1066P^*. Sanger sequencing of the *rpoB* and *rpoC* genes further revealed that the three remaining halo-producing non-*prfA* suppressors also carried point mutations in *rpoC*, namely *rpoC^V1060G^*, which was found twice, and *rpoC^S342F^*. To determine whether mutations in *rpoB* and *rpoC* affect the sugar-dependent PrfA inhibition, PlcB assays were performed with the strains carrying the *rpoB^S1128F^* and *rpoC^R328H^* mutation. With the exception of the *prfA* deletion mutant, all tested strains produced a white halo on LB agar plates supplemented with 0.2% activated charcoal, when no additional carbon source was added. In contrast, only the two strains with mutations in *rpoB* and *rpoC* showed PlcB activity in the presence of glucose or cellobiose thus indicating that PrfA is still active under this condition in these strains (Fig. S1A). Previous studies indicated that alterations in growth and sugar consumption rates can already affect the degree of sugar-dependent PrfA repression [33,34]. Therefore, growth assays were performed revealing that the mutations in *rpoB* and *rpoC* already lead to a minor growth defect in BHI broth. Interestingly, strains pIMK2-*P_actA_-cat rpoB^S1128F^* and pIMK2-*P_actA_-cat rpoC^R328H^* were nearly unable to grow in LSM supplemented with either glucose or cellobiose as sole carbon source. The standard deviation at later time points indicates that both strains formed suppressors (Fig. S1B). As the relief of sugar-dependent PrfA inhibition is likely caused by reduced consumption of glucose and cellobiose, *rpoB* and *rpoC* suppressors were not further studied.

The newly identified PrfA mutations are distributed across the entire protein and map to 13 novel amino acid positions (Fig. 1). The D33A and Q35K/Q35R mutations target residues located in a loop between the first and second β-strand of PrfA. F29, which has been previously already identified, is located in the same loop. Amino acids K64, V87 and V96, which were substituted with glutamate, isoleucine and phenylalanine or leucine, respectively, are situated on β-strands 4, 6 and 7. K64 is part of the glutathione binding site [18,20], while V87 and V96 are surface exposed. Tyrosine on position 11, which is located within the first α-helix, was exchanged for aspartate. Amino acids V124, located near the dimerization interface, and Y126, which is involved in glutathione binding [18,20], were substituted with alanine and phenylalanine, respectively. Residues L148 and V153 on α-helix 4 were mutated to phenylalanine and methionine, respectively. D222, positioned within α-helix 8, was replaced by glutamate and aspartate at position 167 lies at the α5-β9 interface and was mutated to tyrosine (Fig. 1).

**Fig. 1:**
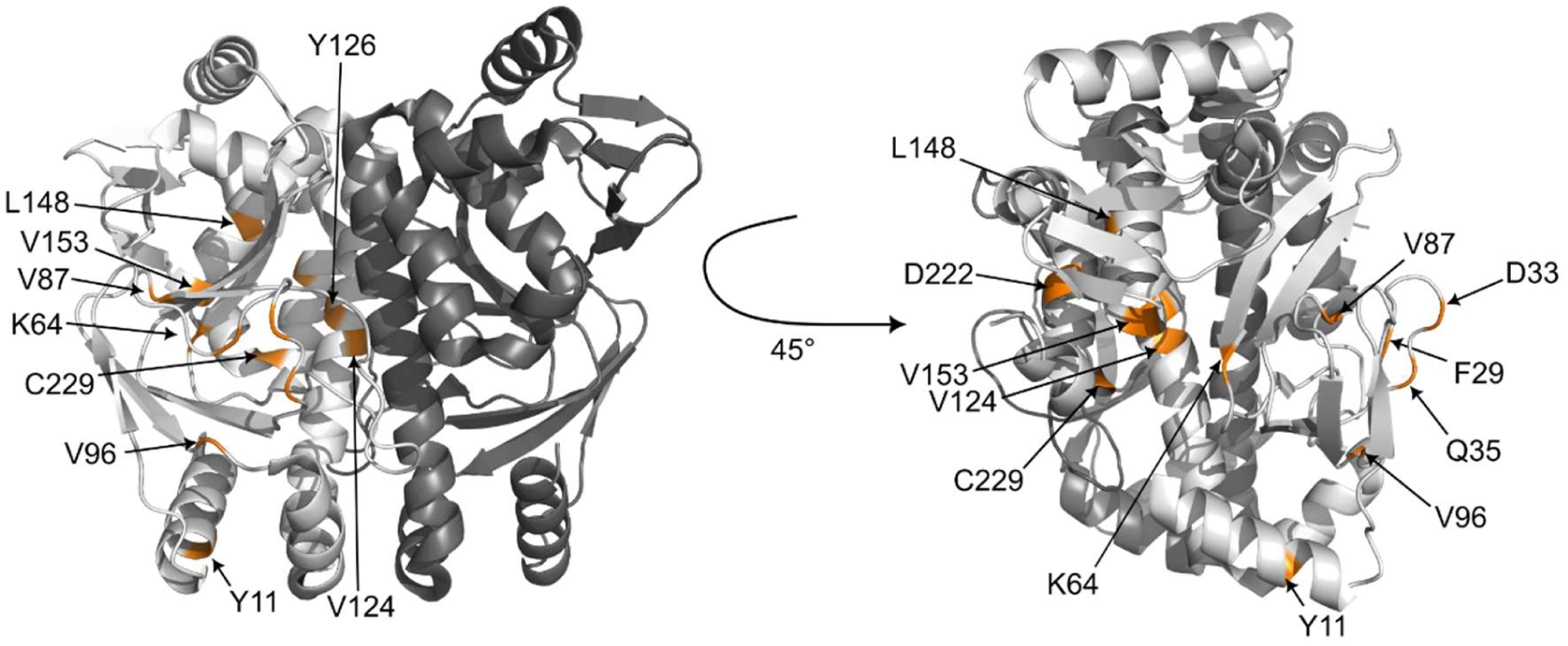
Localization of PrfA point mutations. The dimeric protein structure of PrfA in its DNA binding conformation (PDB: 5LRS) [20] was modified using the PyMOL Molecular Graphics System (Version 3.1.8, Schrödinger, LLC). The individual monomers of PrfA are depicted in light and dark grey. The identified mutations are highlighted in orange.

To assess the impact of these newly identified mutations on PrfA activity, *prfA* alleles carrying individual point mutations were reintroduced into the native locus in a *prfA* deletion strain. For amino acid positions, where multiple substitutions were identified, one was selected for further analysis. A strain, where the wild type *prfA* allele was reinserted into the native locus was used as a control and is further referred to as *prfA^wt^*. Additionally, a strain that produces the previously characterized hyperactive variant PrfA^G145S^ was constructed as reference [14,29].

### Only *prfA^L148F^* has a growth defect in LSM glucose

Constant virulence gene expression is an energetic burden for *L. monocytogenes*. The production of the hyperactive PrfA variant PrfA^G145S^ therefore leads to a growth deficit in both complex and minimal media [14,15,35]. Growth assays in LSM glucose revealed that only *L. monocytogenes* strains harboring the *prfA^G145S^* and *prfA^L148F^* alleles grew slightly slower compared to the wild type strain (Fig. 2), while all other strains behaved like wild type (Fig. S2).

**Fig. 2:**
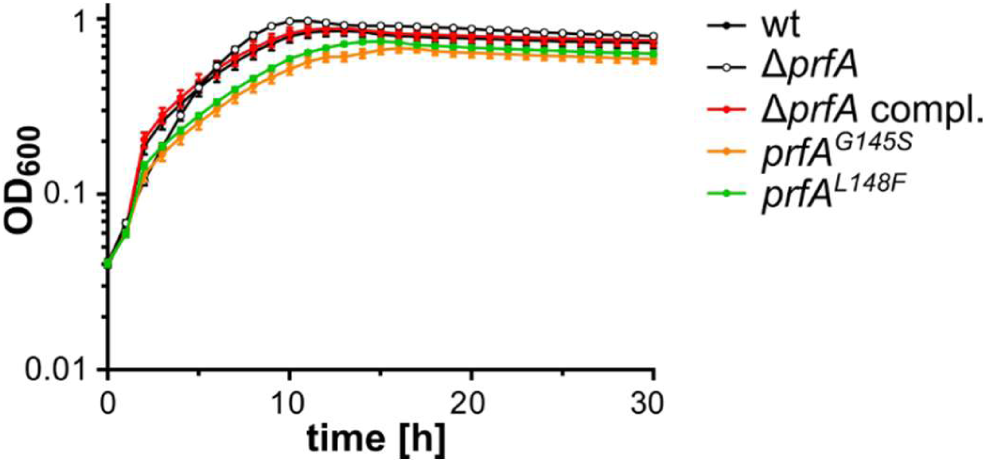
Growth of EGD-e *prfA^G145S^* and EGD-e *prfA^L148F^* in LSM glucose. Overnight cultures of the indicated *L. monocytogenes* strains were diluted in fresh BHI and grown until an OD_600_ of 0.3. 2 ml of the cultures were harvested, washed twice in LSM glucose and resuspended in 1 ml LSM glucose. The cell suspension was adjusted to an OD_600_ of 0.1 and transferred into wells of a 96-well plate. Growth of the strains was measured every 15 min for 30 hours. Averages and standard deviations of three independent experiments are plotted.

### Diverse growth phenotypes of PrfA* variants in LSM glucose-1-phosphate

Within the host, *L. monocytogenes* uses mainly glycerol and hexose phosphates as carbon sources [11,36,37]. The growth on hexose phosphates such as glucose-1-phosphate depends on the activity of the hexose phosphate transporter Hpt, whose expression is under the control of PrfA [11,12]. We have previously shown that the *L. monocytogenes* wild type strain is unable to grow on LSM with glucose-1-phosphate as sole carbon sources when it was pre-grown in BHI, a condition where PrfA is inactive [38]. Here, we tested the ability of the newly identified PrfA* variants to grow in LSM glucose-1-phosphate (Fig. 3). *L. monocytogenes* strain *prfA^G145S^,* which served as a positive control, grows rapidly in this medium. A similar growth phenotype was observed for *L. monocytogenes* strains harboring the *prfA^F29L^, prfA^L148F^, prfA^Y126F^* and *prfA^V96F^*alleles. In contrast, growth of strains *prfA^V124A^, prfA^D222E^, prfA^V87I^, prfA^D33A^, prfA^K64E^, prfA^C229R^, prfA^Y11D^* and *prfA^V153M^* was reduced compared to *prfA^G145S^*. Surprisingly, strain *prfA^Q35K^*displayed an accelerated growth compared to the positive control, while *prfA^D167Y^*behaved like the *L. monocytogenes* wild type strain (Fig. 3).

**Fig. 3:**
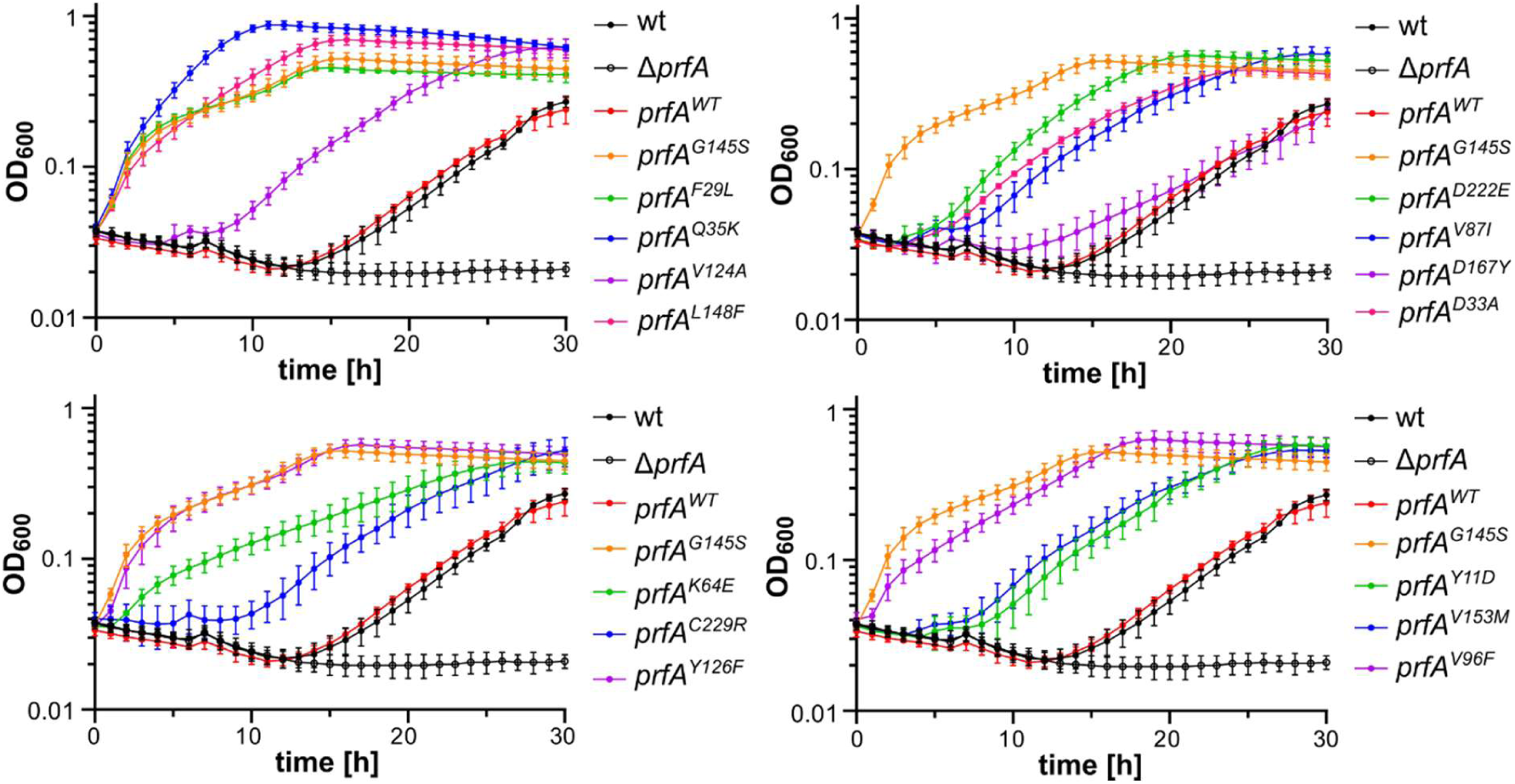
Effect of PrfA mutations on growth in LSM glucose-1-phosphate. Overnight cultures of the indicated *L. monocytogenes* strains were diluted in fresh BHI and grown until an OD_600_ of 0.3. 2 ml of the cultures were harvested, washed twice in LSM glucose-1-phosphate and resuspended in 1 ml LSM glucose-1-phosphate. The cell suspension was adjusted to an OD_600_ of 0.1 and transferred into wells of a 96-well plate. Growth of the strains was measured every 15 min for 30 hours. Averages and standard deviations of three independent experiments are plotted.

### Temperature- and sugar-dependent regulation of PrfA* variants

The translation of the PrfA protein is prevented at low temperatures by an mRNA-thermometer [16]. To assess whether the PrfA translation, and therefore it’s activity, is still dependent on temperature, PlcB assays were performed. At 37°C, halo formation could be observed for all tested strains except for the *prfA* deletion strain. When the PlcB assay plates were incubated at 30°C, no halos were formed by the *L. monocytogenes* wild type strain and the *prfA* complementation strain. In contrast, no temperature-dependence could be observed for any of the PrfA* variants as all strains formed white halos (Fig. 4). In addition to the temperature-dependence, PlcB assays were used to assess the sugar-dependent inhibition of PrfA and PrfA* variants. The *L. monocytogenes* wild type strain and the *prfA* complementation strain did not produce halos in the presence of glucose or cellobiose (Fig. 4), consistent with previous findings [32]. In contrast, all *L. monocytogenes* strains harboring *prfA\** alleles formed halos to varying extents. Interestingly, PlcB and thus PrfA were still active in most of the *L. monocytogenes* strains harboring *prfA\** alleles, when they were grown in the presence of glucose or cellobiose at 30°C. This was especially pronounced for strains *prfA^F29L^, prfA^Q35K^, prfA^V96F^, prfA^Y126F^, prfA^G145S^, prfA^L148F^* and *prfA^D222E^* (Fig. 4).

**Fig. 4:**
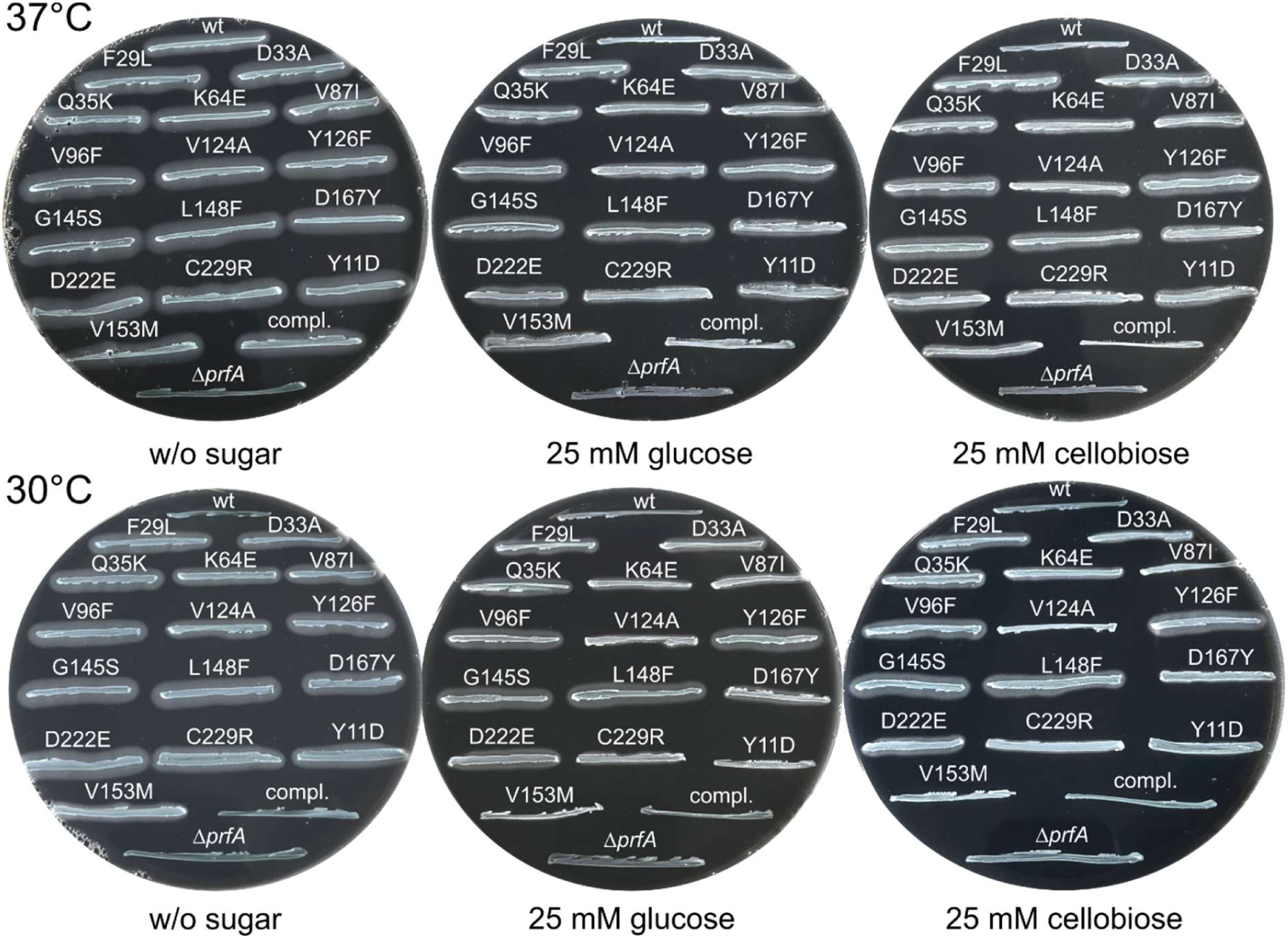
Lecithinase activity assay for assessing temperature- and sugar-dependence of PrfA activity. The indicated PrfA* variants were streaked onto LB agar containing activated charcoal, egg yolk, and where indicated, 25 mM glucose or cellobiose. The *L. monocytogenes* wild type strain EGD-e (wt), the *prfA* deletion strain and the *prfA* complementation strain (compl.) were used as reference. The plates were incubated for a day at 37°C (upper panel) or 30°C (lower panel). Representative images of at least three independent experiments are shown.

PlcB assays only serve as a qualitative read-out of PrfA activity. To determine the impact of sugars on PrfA activity in a quantitative manner, promoter-activity assays were performed. For this purpose, a *P_actA_-lacZ* promoter fusion was integrated into the *L. monocytogenes* wild type strain as well as *L. monocytogenes* strains harboring the indicated *prfA\** alleles. The resulting strains were subsequently grown in either LSM glucose or LSM cellobiose, the latter of which leads to a stronger PrfA inhibition (Fig. 5). The *actA* promoter activity was barely affected by the sugars in strains *prfA^G145S^, prfA^F29L^, prfA^Q35K^, prfA^V124A^, prfA^V87I^, prfA^K64E^, prfA^C229R^, prfA^Y11D^, prfA^V153M^* and *prfA^V96F^*. In contrast, strong cellobiose-dependent PrfA inhibition was observed for strains *prfA^L148F^*, *prfA^D167Y^* and *prfA^D33A^* (Fig. 5). These findings already clearly show that the impact of the PrfA mutations can differ between PrfA-regulated genes, as strong PlcB activity was observed for *prfA^L148F^*and *prfA^D167Y^* in the presence of cellobiose (Fig. 4), whereas *actA* expression was significantly inhibited by cellobiose (Fig. 5).

**Fig. 5:**
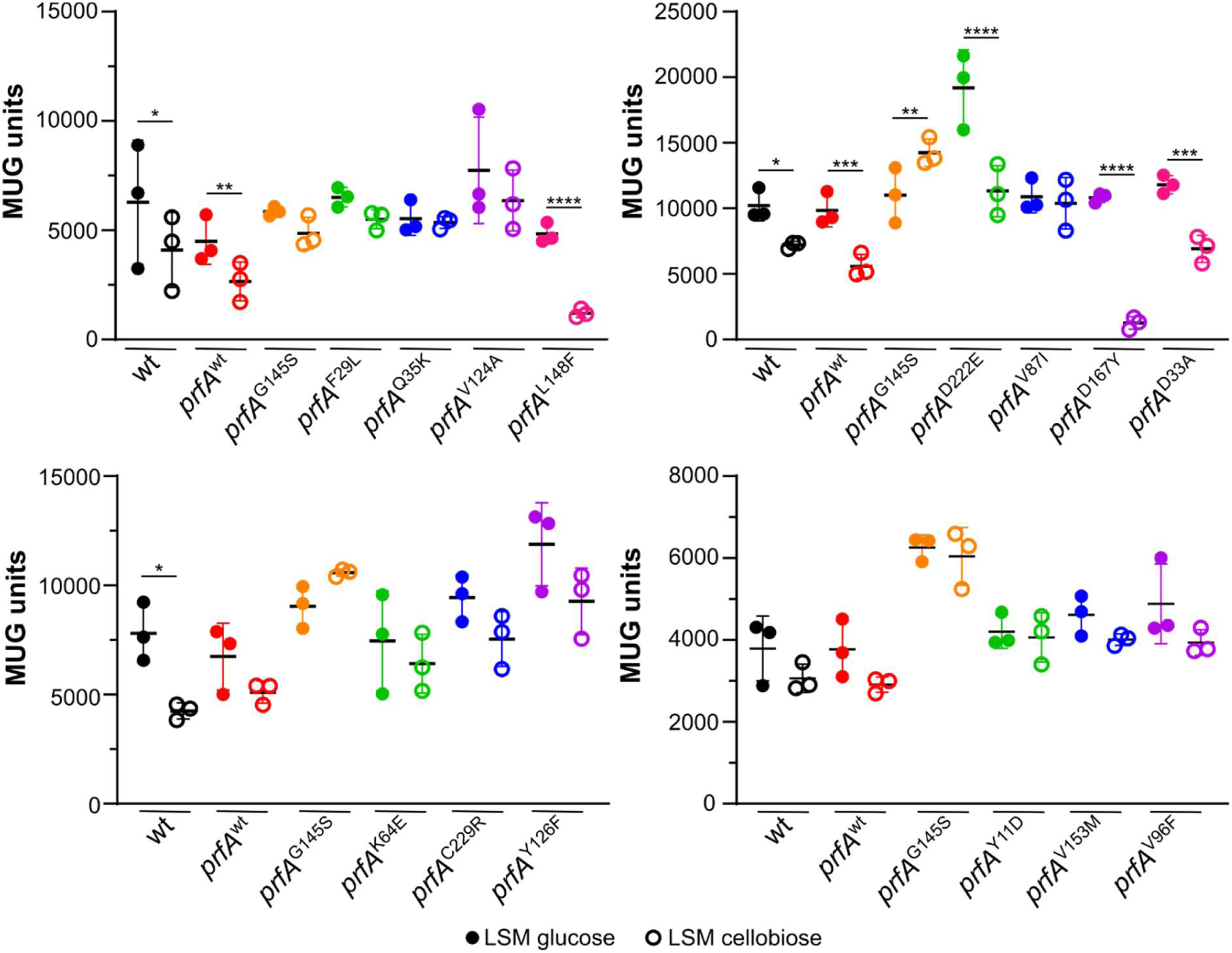
Promoter activity assays to assess sugar-dependent inhibition of PrfA. The activity of the *actA* promoter was determined for the *L. monocytogenes* wild type EGD-e and *L. monocytogenes* strains, in which the wild type or mutated *prfA* gene was re-introduced into the native *prfA* locus. The strains were grown in either LSM glucose (filled circles) or LSM cellobiose (open circles). Averages and standard deviations of three independent experiments are plotted. For statistical analysis, an ordinary two-way ANOVA with Šídák’s multiple comparison test was performed (*, *p* < 0.05; **, *p* < 0.01; ***, *p* < 0.001; ****, *p* < 0.0001).

### Glutathione-dependency of PrfA* variants

The activity of PrfA depends on its cofactor glutathione, which is synthesized by the glutathione synthetase GshF in *L. monocytogenes* [39]. Previous studies already showed that this dependency is lost in PrfA^G145S^ [19]. To assess the glutathione-dependency of the newly identified PrfA* variants, *actA* promoter assays were performed with strains harboring the *prfA\** alleles in either wild type or Δ*gshF* background. In addition to PrfA^G145S^, PrfA* variants with the amino acid substitutions Y126F and V153M were fully active in the absence of glutathione. Intermediate glutathione dependency was observed for PrfA^F29L^, *prfA^Q35K^*, PrfA^V124A^, PrfA^L148F^, PrfA^D222E^, PrfA^D33A^, PrfA^K64E^, PrfA^Y11D^ and PrfA^V96F^. PrfA* variants PrfA^V87I^, PrfA^D167Y^ and PrfA^C229R^ were still fully dependent on glutathione (Fig. 6).

**Fig. 6:**
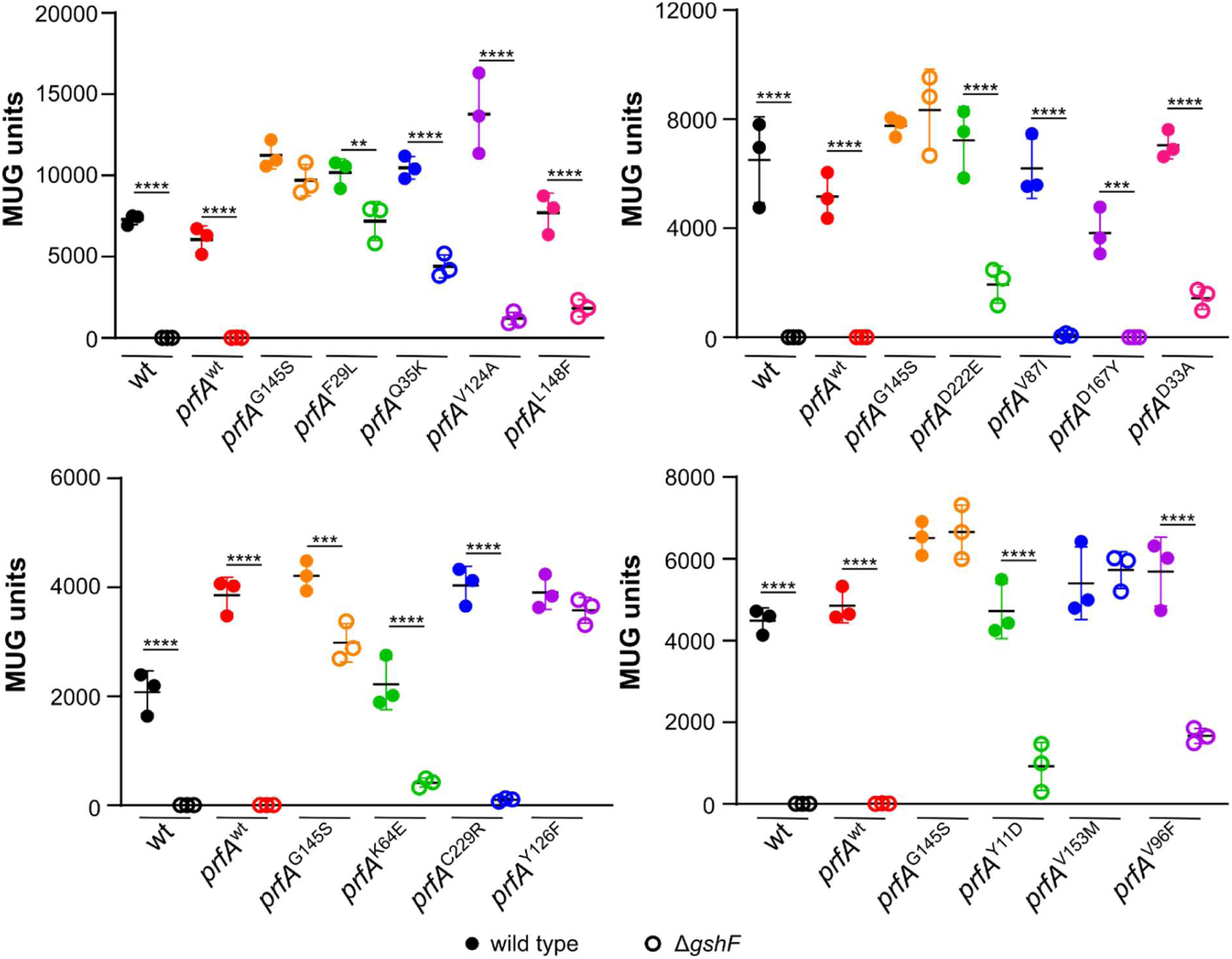
Promoter activity assays to assess glutathione-dependency of PrfA. The activity of the *actA* promoter was determined for the *L. monocytogenes* wild type EGD-e and *L. monocytogenes* strains, in which the wild type or mutated *prfA* gene was re-introduced into the native *prfA* locus. The strains were either able to produce glutathione (filled circles) or lack the glutathione synthetase GshF and are thus unable to produce glutathione (open circles). Averages and standard deviations of three independent experiments are plotted. For statistical analysis, an ordinary two-way ANOVA with Šídák’s multiple comparison test was performed (**, *p* < 0.01; ***, *p* < 0.001; ****, *p* < 0.0001).

### Impact of PrfA* mutations on intracellular replication

Based on their phenotypes (summarized in Table S4), a subset of strains harboring *prfA\** alleles were selected for intracellular replication assays with J774A.1 murine macrophages. Strain *prfA^V124A^* was chosen, because it showed a slightly higher *actA* promoter activity in LSM glucose compared to *prfA^G145S^* (Fig. 5 and 6). Moreover, amino acid V124 is located at the dimerization interface (Fig. 1) and substitution could potentially affect dimerization and thus activity of PrfA. The intracellular growth of strain *prfA^L148F^* was assessed as it was the only PrfA* strain displaying a growth defect in LSM glucose (Fig. 2). Strain *prfA^D222E^* was selected, because it showed a comparable *actA* promoter activity to *prfA^G145S^*, however, it could still be partially inhibited by cellobiose and was partially glutathione-dependent (Fig. 5 and 6). Furthermore, strain *prfA^D167Y^*was analyzed due to its wild type-like growth in LSM supplemented with glucose-1-phosphate (Fig. 3). Lastly, strain *prfA^V87I^* was chosen for intracellular growth assays because the *actA* promoter activity was not affected in the presence of cellobiose, while the PrfA variant was fully glutathione-dependent (Fig. 5 and 6). *L. monocytogenes* strain *prfA^wt^*, in which the wild type *prfA* gene was reinserted, served as wild type control, since it was constructed in the same manner as the strains harboring the *prfA\** alleles. Strain *prfA^G145S^* showed a slightly enhanced intracellular replication compared to *prfA^wt^*(Fig. 7). Surprisingly, a slightly decreased intracellular replication relative to *prfA^wt^*was observed for *prfA^D222E^*, *prfA^V124A^* and *prfA^L148F^*. An even more pronounced reduction in intracellular replication was measured for strains *prfA^V87I^* and *prfA^D167Y^* (Fig. 7). These results suggest that enhanced PrfA activity in *in vitro* assays, does not necessarily translate into an enhanced virulence potential.

**Fig. 7:**
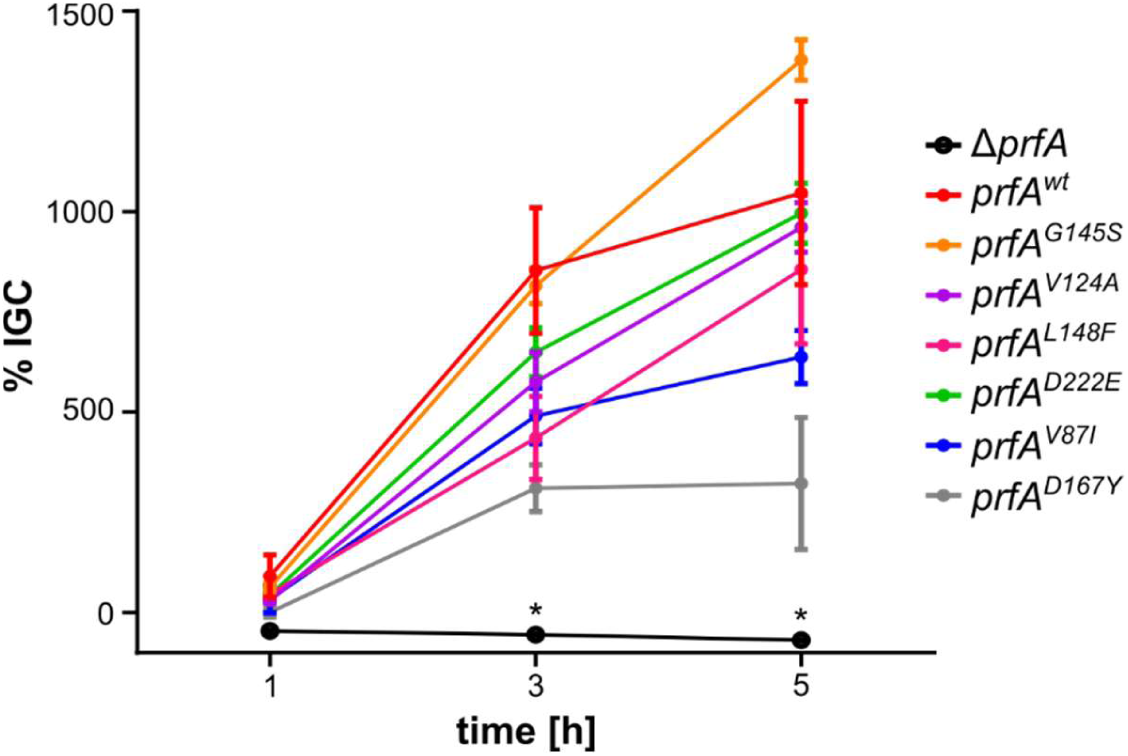
Impact of PrfA mutations on intracellular replication. J774A.1 cells were infected at an MOI of 1 with the indicated *L. monocytogenes* strains. The number of intracellular bacteria was determined after 1, 3 and 5 hours and the intracellular growth coefficient (% IGC) was calculated as described in the methods section. Averages and standard deviations of a representative experiment performed with biological triplicates is shown. For statistical analysis, a repeated-measures two-way ANNOVA with Dunnett’s multiple comparison test was performed in GraphPad Prism to compare the *L. monocytogenes prfA^wt^* strain with the other tested strains (*, *p* < 0.05).

### C38 does not seem to be involved in sugar-dependent PrfA inhibition

A previous study suggested that the cysteine at position 38 of PrfA is potentially phosphorylated by the EIIA/EIIB component ManL in response to glucose, thereby rendering it inactive [40]. Surprisingly, no suppressors with alterations of C38 of PrfA were isolated in our screens. To determine the impact of phospho-mimetic and phospho-ablative mutations on sugar-dependent inhibition, the *prfA* alleles *prfA^C38E^*, *prfA^C38A^*, and *prfA^C38S^* were reintroduced into the native *prfA* locus. In accordance with the previous study [40], PlcB assays with these strains revealed that PrfA^C38E^ appeared to be inactive as no halo formation could be observed for any of the tested conditions. In contrast, strains harboring the *prfA^C38A^* and *prfA^C38S^* allele were still able to produce halos in the absence of sugars; however, both PrfA variants were still subject to sugar-dependent inhibition as halo formation is lost upon addition of glucose or cellobiose (Fig. 8A). Promoter activity assays further revealed that the exchange of cysteine to alanine or serine at position 38 slightly reduced the activity of PrfA. Furthermore, the activity of the two PrfA variants PrfA^C38A^ and PrfA^C38S^ was still fully dependent on the presence of glutathione (Fig. 8B). Consistent with the PlcB assays, the exchange to glutamic acid led to complete loss of PrfA activity irrespective of the absence or presence of the cofactor glutathione (Fig. 8B). These results suggest that mutations of cysteine at position 38 affect PrfA activity in general, but not specifically the sugar-dependent PrfA inhibition.

**Fig. 8:**
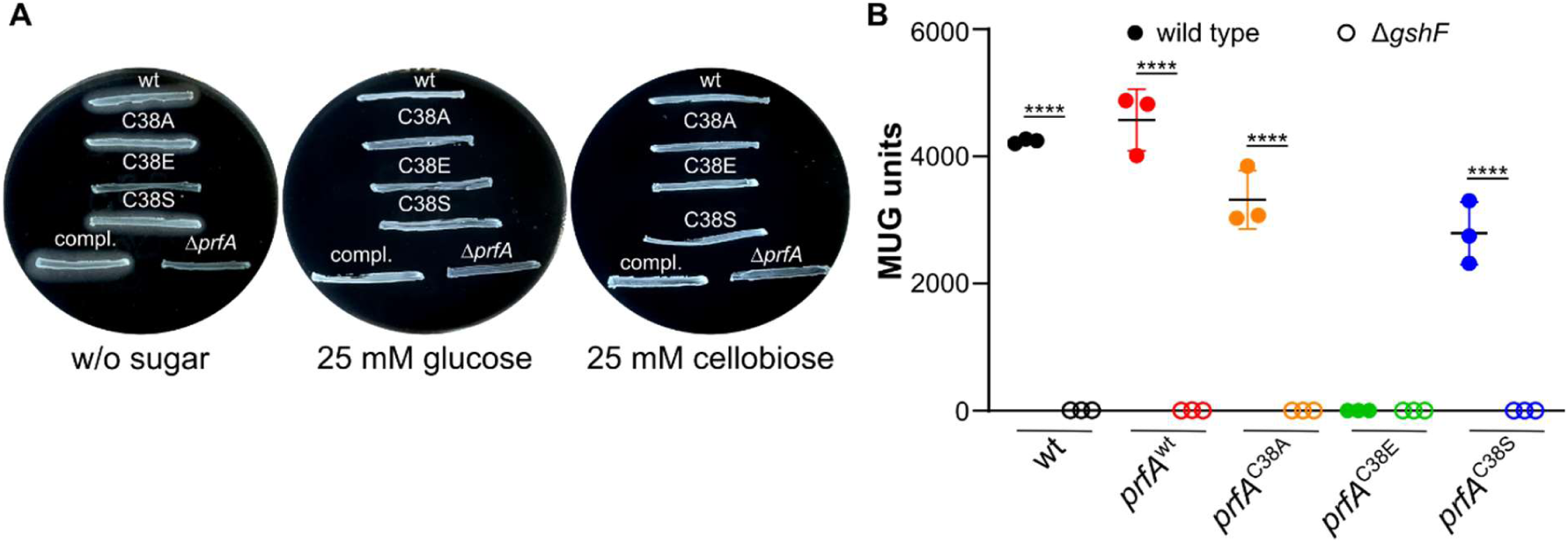
(A) PlcB Assay. Strains producing C38 variants of PrfA were streaked onto LB agar containing activated charcoal, egg yolk, and where indicated, 25 mM glucose or cellobiose. The *L. monocytogenes* wild type strain EGD-e (wt), the *prfA* deletion strain and the *prfA* complementation strain (compl.) were used as reference. The plates were incubated for a day at 37°C and representative images of three independent experiments are shown. (B) Promoter activity assay. The activity of the *actA* promoter was determined for the *L. monocytogenes* wild type EGD-e and *L. monocytogenes* strains, in which the wild type or mutated *prfA* gene was re-introduced into the native *prfA* locus. The strains were either able to produce glutathione (filled circles) or lack the glutathione synthetase GshF and are thus unable to produce glutathione (open circles). Averages and standard deviations of three independent experiments are plotted. For statistical analysis, an ordinary two-way ANOVA with Šídák’s multiple comparison test was performed (****, *p* < 0.0001).

## Discussion

In an attempt to identify factors involved in the sugar-dependent PrfA inhibition, a suppressor screen was performed. For this, *L. monocytogenes* strains carrying a chloramphenicol resistance cassette, which is under the control of either the *actA* or *hly* promoter, were streaked on agar plates containing cellobiose and chloramphenicol. The plates additionally contained egg yolk and activated charcoal to only select colonies that possess PlcB activity and are thus able to grow due to active PrfA. Sequencing of the *prfA* gene revealed that 56 of the 64 isolated suppressors acquired mutations rendering PrfA hyperactive. Similar results were obtained in a previous study, where a transposon-based screening was used to identify genes involved in the sugar-dependent PrfA inhibition. For this purpose, transposon mutants with a PrfA-activated phenotype were isolated and more than 80% of these strains carried mutations in *prfA* [27]. In the past years, at least 11 amino acid positions whose substitution results in a PrfA* phenotype have been identified and characterized to a varying extent, such as I45S, A94T, L140F, G145S, G155S and P219S [26–32,41,42]. Astonishingly, 13 novel PrfA* variants at amino acid positions that have not previously been associated with hyperactivity were identified in the here conducted suppressor screen. PlcB activity, growth, and promoter activity assays were used to assess the effects of individual PrfA mutations on PrfA activity in response to temperature and sugar availability, as well as to determine whether the PrfA* variants remained dependent on glutathione as a cofactor (summarized in Table S4). Interestingly, all strains producing PrfA* variants exhibited PlcB activity at 30°C, suggesting that the mutations reduce the impact of temperature-dependent post-transcriptional regulation of PrfA. Consistent with this, PlcB activity was also detected for PrfA* strains harboring the amino acid substitutions G145S, I45S and L140F grown at 20°C [26].

The results of the PlcB activity, growth, promoter activity and virulence assays indicate that the effects of PrfA* mutations are both variant- and gene-dependent. The level of activation varied between PrfA* variants and differed also for the genes that are controlled by PrfA. For instance, high *actA* promoter activity levels were measured for the *L. monocytogenes* strain producing PrfA^V124A^, which exceeded the ones of PrfA^G145S^, while the PrfA^V124A^ strain only showed moderate growth on glucose-1-phosphate and reduced intracellular growth compared to PrfA^G145S^. Interestingly, a strain carrying the D167Y substitution in PrfA showed PlcB activity even in the presence of sugars or low temperatures, suggesting that this mutation leads to a PrfA* phenotype. On the contrary, in promoter activity and growth assays performed with glucose-1-phosphate, this strain behaved similar to the wild type strain. Moreover, in infection assays, this strain even exhibited reduced intracellular growth compared to the wild type. These examples illustrate that one cannot predict virulence potential of PrfA* strains simply by assessing the expression or activity of single PrfA-dependent genes. Varying levels of PrfA activation due to PrfA* mutations have already been described previously [14,27,41].

Crystal structure analysis of homodimers of PrfA* variants revealed the existence of three forms of PrfA: 1) the inactive form, e.g. the PrfA wild type lacking its allosteric cofactor glutathione, 2) an intermediately active form, which was observed for PrfA^L140H^, PrfA^L140F^ and PrfA^A218G^ and 3) an active form, which was observed for PrfA^G145S^ and the PrfA^wt^-glutathione complex [20,27,43]. In the active PrfA dimer, both HTH motifs are folded and able to interact with DNA. In PrfA^wt^, this is only achieved when all three residues of glutathione are correctly positioned within the binding site. Particularly important are the interactions of the carboxyl group of glycine of glutathione with amino acid positions Y126, Y154 and S177 of PrfA. Correct binding of glutathione leads to the collapse of the intraprotein tunnel harboring the binding site, which must occur in both PrfA monomers to form the fully active form of the homodimer [20]. Thus, glutathione shifts PrfA toward its active, DNA-binding conformation by stabilizing the HTH motif. The hyperactivating mutation G145S shifts PrfA permanently toward the active conformation and thereby bypasses the requirement of glutathione binding [19,20]. Promoter activity assays revealed that PrfA* variants with mutations Y126F and V153M are also active in the absence of glutathione. The substitution of tyrosine at position 126 with phenylalanine is considered conservative and is not expected to substantially alter the hydrophobic environment of the intraprotein tunnel. However, it removes the hydroxyl group of Y126, which participates in interactions with the carboxyl group of glycine of glutathione [20]. Given that PrfA^Y126F^ retains activity in the absence of glutathione, one possible explanation is that the mutation alters the conformational equilibrium of PrfA, favoring the active state over the inactive state. Surprisingly, AlphaFold3 [44,45] predictions of the PrfA^Y126F^ dimer suggest that the overall structure remains more similar to the inactive conformation of PrfA (Fig. S3). However, AlphaFold3, due to its template bias, may not be able to fully predict how a single amino acid substitution affects protein stability, conformational dynamics or the balance between different conformational states [46]. In the second glutathione-independent PrfA* variant, valine 153 is substituted with methionine. Valine at position 153 forms a hydrogen bond with lysine 64 of PrfA, which is required for glutathione-mediated activation [20,47]. Methionine has a larger and more flexible side chain than valine. The V153M substitution could therefore alter local packing within the intraprotein tunnel and consequently influence the conformational equilibrium of PrfA by favoring its active conformation. Several PrfA* variants exhibiting partial glutathione dependency were identified such as PrfA^V124A^, PrfA^L148F^, PrfA^D222E^ or PrfA^V96F^. It is tempting to speculate that these variants may adopt an intermediate active fold, similar to that described for PrfA^L140H^, PrfA^L140F^ and PrfA^A218G^. This intermediate active conformation of the PrfA dimer is characterized by one HTH motif adopting the folded, active conformation, while the other remains in the non-active conformation of PrfA^wt^ [27]. Taken together, these observations suggest that different PrfA* mutations may influence the equilibrium between active and inactive conformations in different ways, which could explain why some variants are more dependent on glutathione-based activation than others.

A previous study proposed that the cysteine at position 38 may be involved in sugar-dependent inhibition of PrfA [40]. Although the PlcB and promoter activity assays performed in this study did not support a role for C38 in sugar-dependent PrfA inhibition, the results indicate that C38 does influence PrfA activity. C38 is located within the N-terminal β-barrel domain, where it is part of a hydrophobic internal core. Substitution of cysteine with alanine, a non-polar and hydrophobic amino acid that is slightly smaller than cysteine, may create a small void within this core. This could reduce local packing and slightly destabilize the protein, resulting in the reduced activity observed in the promoter activity assays. The PrfA^C38S^ variant also showed reduced activity in the promoter activity assays. In this case, replacing cysteine with serine introduces a polar hydroxyl group into the hydrophobic core. This change is energetically less favorable and may disrupt local interactions within this core, thereby reducing the stability of the active conformation of PrfA. However, because serine is similar in size to cysteine, the overall protein structure is likely to remain intact, which could explain the partial reduction in activity rather than a complete loss of function. In contrast, substitution of C38 with glutamic acid introduces a larger, negatively charged residue into the hydrophobic core. This is likely to cause a much greater disruption of the local environment and could compromise the stability of the protein, potentially resulting in misfolding or loss of the functional conformation. This could explain the complete loss of PrfA activity observed for PrfA^C38E^ in both the PlcB and promoter activity assays. However, structural and biochemical experiments are required to determine whether these proposed effects on protein stability and conformation are responsible for the observed changes in PrfA activity.

Taken together, this study in conjunction with the already existing literature shows that PrfA activity can be easily enhanced by substitution of only a single amino acid. However, enhanced *in vitro* activity does not necessarily translate into an increased virulence potential of *L. monocytogenes* PrfA* strains *in vivo*. This highlights once again the multifactorial complexity of PrfA regulation in response to environmental and host cues.

## Materials and methods

### Bacterial strains and growth conditions

All strains and plasmids used in this study are listed in Table S1. *Escherichia coli* strains were cultivated in lysogeny broth (LB) medium *and Listeria monocytogenes* strains in brain heart infusion (BHI) medium or *Listeria* synthetic medium (LSM) at 37°C unless otherwise stated. LSM was prepared as previously described using glucose or cellobiose as sole carbon source [38]. For the preparation of LSM, a complete amino acid mix containing low concentrations of branched chain amino acids (BCAA), namely 80 µM L-isoleucine, 80 µM L-leucine and 90 µM L-valine, was used. When required, media were supplemented with antibiotics at the following concentrations: for *E. coli* cultures: 100 µg ml^-1^ ampicillin (amp), 30 µg ml^-1^ chloramphenicol (cat), 50 µg ml^-1^ kanamycin (kan); for *L. monocytogenes* cultures: 7.5 µg ml^-1^ chloramphenicol, 50 µg ml^-1^ kanamycin, 5 µg ml^-1^ erythromycin (ery), 30 µg ml^-1^ nalidixic acid (nal).

### Strain and plasmid construction

All plasmids used in this study are listed in Table S2. Plasmids were constructed for the suppressor screens, in which the expression of a chloramphenicol resistance cassette is driven by the *actA* or *hly* promoter. For this purpose, *P_actA_* and *P_hly_* were amplified from genomic DNA of the *L. monocytogenes* wild type strain EGD-e by PCR using primer pairs JR295/JR296 and JR299/JR300, respectively. The chloramphenicol resistance gene (*cat*) was amplified from plasmid pPL2 using primer pairs JR297/JR298 (*P_actA_*) and JR301/JR298 (*P_hly_*). The *P_actA_* and *P_hly_* fragments were subsequently fused with the corresponding *cat* gene using primers JR295/JR298 and JR299/JR298, respectively. The promoter-*cat* constructs were digested with *EcoR*I and ligated into plasmid pIMK2, which had been cut with the same enzyme. The resulting plasmids pIMK2-*P_actA_-cat* and pIMK2-*P_hly_-cat* were recovered in *E. coli* DH5α yielding strains EJR278 and EJR279, respectively. Both plasmids were subsequently transformed into *L. monocytogenes* via electroporation yielding strains EGD-e pIMK2-*P_actA_-cat* (LJR332) and EGD-e pIMK2-*P_hly_-cat* (LJR333).

For the reinsertion of *prfA* variants into the *prfA* deletion strain, multiple pKSV7-*prfA* plasmids were constructed. For this purpose, a fragment consisting of the *prfA* gene and 1-kb flanking regions were amplified from genomic DNA of the *L. monocytogenes* wild type strain EGD-e or of isolated suppressors carrying mutations in *prfA* by PCR using primer pair JR258/JR261. The resulting fragments were digested with *Kpn*I/*Sal*I and ligated into plasmid pKSV7 that had been cut with the same enzymes. The pKSV7-*prfA* plasmids were recovered in *E. coli* XL10 Gold and subsequently electroporated in the *L. monocytogenes prfA* deletion strain (BUG2214). The *prfA* gene was reinserted into Δ*prfA* by allelic exchange as previously described [48]. The reinsertion of *prfA* was confirmed via lecithinase activity assays and PCR.

For the isolation of strains EGD-e *prfA^F29L^* (LJR381) and EGD-e *prfA^Q35K^* (LJR383), log-phase cultures of the *L. monocytogenes* wild type EGD-e were grown in BHI broth. Subsequently, 1 ml of culture was harvested, washed in LSM containing glucose-1-phosphate or glucose-6-phosphate as sole carbon source and used for growth experiments in the respective LSM using a plate reader. Under this condition, *prfA* is barely active and the wild type strain is unable to grow as previously shown [38]. Suppressors could be isolated after prolonged incubation and sanger sequencing revealed point mutations within the *prfA* gene.

For markerless deletion of *gshF*, 1 kb DNA fragments up- and downstream of *gshF* were amplified using primer pairs JR319/JR320 and JR321/JR322, respectively. The resulting PCR products were fused in a second PCR using primers JR319/JR322. The deletion cassette was digested with *Sal*I/*Kpn*I and ligated into plasmid pKSV7 that had been cut with the same enzymes. pKSV7-Δ*gshF* was recovered in *E. coli* XL10 Gold yielding strain EJR276. Plasmid pKSV7-Δ*gshF* was subsequently electroporated into the *L. monocytogenes* wild type strain EGD-e and *gshF* deleted by allelic exchanges as previously described yielding strain EGD-e Δ*gshF* (LJR350). The *gshF* gene was deleted in a similar manner in the strains, in which wild type or mutated variants of *prfA* were reinserted. Deletion of *gshF* was confirmed by PCR.

For promoter activity assays, plasmid pPL3e-*P_actA_-lacZ* was isolated from EJR286 [33] and transformed into *E. coli* S17-1 yielding strain EJR331. Strain EJR331 was subsequently used as a donor to transfer plasmid pPL3e-*P_actA_-lacZ* by conjugation into *L. monocytogenes* strains, in which wild type or mutated variants of *prfA* were reinserted and the strains, in which *gshF* had been deleted as previously described [49].

### Lecithinase (PlcB) activity assay

For a qualitative PlcB activity assay, which was performed as previously described with minor modifications [50], single colonies of *L. monocytogenes* strains were streaked on LB agar plates supplemented with 0.2% activated charcoal and 2% of an egg yolk suspension, prepared by mixing one egg yolk with one volume of 1x phosphate-buffered saline (PBS, pH 7.4). For sugar-dependent PrfA inhibition, 25 mM of glucose or cellobiose were added to the plates. Inoculated plates were incubated for a day at 37°C and subsequently photographed.

### Suppressor screen

*L. monocytogenes* strains containing pIMK2-*P_actA_-cat* and pIMK2-*P_hly_-cat* (LJR332 and LJR333) were grown overnight in BHI medium supplemented with kanamycin. The suppressor screen was performed directly with overnight cultures and with log-phase cultures. For the preparation of log-phase cultures, overnight cultures were diluted in fresh BHI medium to an OD_600_ of 0.1 and grown at 37°C and shaking until they reached an OD_600_ of 0.6. Then, 100 µl of overnight and log-phase cultures that were adjusted to an OD_600_ of 0.1 were plated on LB plates containing 0.2% activated charcoal, 2% egg yolk suspension, 25 mM cellobiose and varying chloramphenicol concentrations (30, 40 or 50 µg ml^-1^). The plates were incubated for up to 4 days at 37°C. *L. monocytogenes* colonies that grew on these plates and possessed lecithinase activity were re-streaked twice on selective plates.

Additionally, suppressors were isolated in LSM containing glucose-1-phosphate or glucose-6-phosphate as sole carbon source, a condition in which the wild type strain EGD-e is unable to grow [38]. For this purpose, EGD-e overnight cultures were diluted to an OD_600_ of 0.1 in fresh BHI broth and grown at 37°C until they reached an OD_600_ of 0.3. Subsequently, 2 ml of the cells were collected, washed twice in 1 ml LSM glucose-1-phosphate and LSM glucose-6-phosphate, respectively, and resuspended in the corresponding medium. The cultures were adjusted to an OD_600_ of 0.1 and 200 µl of the cell suspension transferred into wells of a 96-well plate. The plate was incubated at 37°C with orbital shaking in an BioTek Epoch 2 microplate reader and OD_600_ values measured every 15 min. Growth on glucose-1-phosphate and glucose-6-phosphate was observed after approximately 24 and 32 hours, respectively. Suppressors were re-streaked twice on LSM glucose-1-phosphate and glucose-6-phosphate agar plates. For both suppressor screens, the *prfA* gene and promoter region were amplified by PCR and mutations identified by Sanger sequencing.

### Whole genome sequencing

Genomic DNA of LJR366, LJR367 and LJR368 was isolated and send to SeqCenter (Pittsburgh, PA, United States) for whole genome sequencing. Short read sequencing (151 bp paired end) was performed on an Illumina NovaSeq X Plus sequencer (San Diego, CA, United States). The obtained reads were mapped to the *L. monocytogenes* EGD-e reference genome (NC_003210) using Geneious prime v.2025.2.1 (Biomatters Ltd., New Zealand). Single nucleotide polymorphisms (SNPs) with a variant frequency of at least 95% and a coverage of more than 100 reads were considered as mutations.

### Growth curves

Overnight cultures of the indicated *L. monocytogenes* strains were diluted to an OD_600_ of 0.1 in fresh BHI broth. After the cultures reached an OD_600_ of 0.3, 2 ml of the cells were collected, washed twice with 1 ml BHI, LSM glucose, LSM cellobiose or LSM glucose-1-phosphate. The cell pellet was resuspended in 1 ml of the corresponding medium and cultures adjusted to an OD_600_ of 0.1. Afterwards, 200 µl of the cell suspension were transferred into wells of a 96-well plate. The plate was incubated at 37°C with orbital shaking in an BioTek Epoch 2 microplate reader and OD_600_ values measured every 15 min for at least 30 hours. Averages and standard deviations of at least three independent growth assays were plotted.

### β-galactosidase assay

The indicated *L. monocytogenes* strains were cultivated in 10 ml LSM glucose containing 5 µg ml^-1^ erythromycin overnight at 37°C and 200 rpm. The next day, cultures were used to inoculate 15 ml fresh LSM glucose or LSM cellobiose to an OD_600_ of 0.1. The cultures were grown at 37°C and 200 rpm until they reached an OD_600_ of 0.6 – 0.8. The final OD_600_ was measured for all samples prior sample collection. Subsequently, 1 ml of each culture was collected, resuspended in 100 µl assay buffer with Triton X-100 (ABT buffer) (60 mM K_2_HPO_4_, 40 mM KH_2_PO_4_, 100 mM NaCl, 0.1% Triton X-100; pH 7), snap-frozen in liquid nitrogen, and stored at -80° until further use. The β-galactosidase assay was performed as described previously [51]. Briefly, samples were thawed, and 10-fold dilutions were prepared in ABT buffer. 50 µl of each dilution were mixed with 10 µl of 0.4 mg ml^-1^ 4-methyl-umbelliferyl-β-D-galactopyranoside (MUG) substrate (Merck, Darmstadt, Germany) that was dissolved in dimethyl sulfoxide. Reactions containing ABT buffer and the substrate were used as blank. After a 60-minute incubation in the dark, 20 µl of each reaction were transferred into the wells of a black 96-well plate containing 180 µl ABT buffer. Fluorescence values were determined using a Synergy Mx microplate reader (BioTek) at 366 nm excitation and 445 nm emission wavelengths. Concentrations from 0.015625 to 4 µM of the fluorescent standard 4-methylumbelliferone were used to obtain a standard curve. β-galactosidase units, or MUG units, were calculated as:

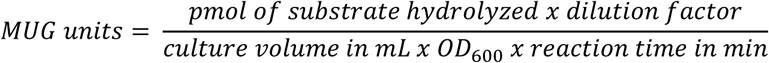

The amount of hydrolyzed substrate was determined from the standard curve as:

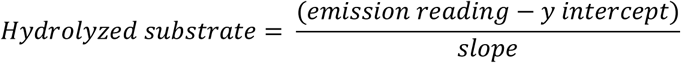

### Mammalian cell cultures and intracellular replication assays

The murine macrophage cell line J774A.1 (ATCC TIB-67^TM^) was used in this study. The J774A.1 cells were grown in Dulbecco’s Modified Eagle Medium (DMEM; Gibco) supplemented with 10% fetal bovine serum (FBS; Gibco) and 1% penicillin and streptomycin (PS; Gibco) under 5% CO_2_ at 37°C. Media change, sub-culturing of J774A.1 cells and intracellular replication assays were carried out as detailed in Borreby et al., [52], with minor modifications. Overnight cultures of the indicated *L. monocytogenes* strains were diluted to an OD_600_ of 0.02 in BHI broth and grown until an OD_600_ of 0.8. J774A.1 cells were infected at a multiplicity of infection (MOI) of 1. Following infection, the cells were incubated for 30 min under 5% CO_2_ at 37°C. After this primary incubation, cells were washed twice with DMEM + 10% FBS and subsequently incubated in DMEM + 10% FBS supplemented with 50 µg/mL gentamicin for an additional 30 min, 2.5 h, or 4.5 h. After the incubation times, cells were washed twice with 500 µl 1x PBS and subsequently lysed using 500 µl 1x PBS + 1% Triton X100. The number of intracellular bacteria was determined by plating serial dilutions on BHI agar. Data from intracellular infection assays were normalized using the intracellular growth coefficient (% IGC), as previously described [28]. % IGC was calculated as:

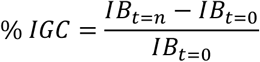

Intracellular replication assays were performed in three biological replicates and were repeated independently three times.

### AlphaFold3 predictions of PrfA^Y126F^ and PrfA^V153M^

Structural models for the homodimers PrfA^Y126F^ and PrfA^V153M^ were generated using AlphaFold3 [44,45]. In total, 250 individual prediction runs were performed for each protein variant. The predicted models were ranked using a composite overall score derived from the calculated pTM and ipTM scores and defined as: Ranking Score = 0.8 ipTM + 0.2·pTM. The top 5 highest-scoring predictions were subsequently superimposed on the crystal structures of apo-PrfA (PDB: 1OMI) and glutathione-bound PrfA (PDB: 5LRR) [20] for structural comparison.

## Acknowledgements

We thank Julia Busse for technical assistance and Nele Sievers for the help with some experiments. We thank Piotr Neumann for valuable and fruitful discussions. We are grateful to Prof. Jörg Stülke for providing JR and MA with laboratory space, equipment and consumables and to the Göttingen Center for Molecular Biosciences (GZMB) and the Dorothea Schlözer Programme for financial support. This work was funded by the German research foundation (DFG) grant RI 2920/3-3 to JR and the Independent Research Fund Denmark grant 4283-00017B to BHK.

