## Supplemental Tables and Figures for "No two are the same: Identification and characterization of novel hyperactive PrfA* variants in *Listeria monocytogenes*"

**Table S1: Bacterial strains used in this study**

| Unique ID | Strain name and resistance | Source |
|---|---|---|
| ***Escherichia coli* strains** | | |
| ANG124 | DH5α pKSV7; $amp^R$ | [1] |
| ANG4242 | XL1-Blue pIMK2; $kan^R$ | [2] |
| ANG5181 | XL1-Blue pPL3e-*lacZ*; $cat^R$ | [3] |
| EJR274 | XL1-Blue pPL3e-$P_{plcA}$-$P_{prfA}$-*prfA*; $cat^R$ | [4] |
| EJR276 | XL10-Gold pKSV7-Δ*gshF*; $amp^R$ | This study |
| EJR278 | DH5α pIMK2-$P_{actA}$-*cat*; $kan^R$ | This study |
| EJR279 | DH5α pIMK2-$P_{hly}$-*cat*; $kan^R$ | This study |
| EJR286 | DH5α pPL3e-$P_{actA}$-*lacZ*; $cat^R$ | [4] |
| EJR297 | XL10-Gold pKSV7-$prfA^{L148F}$; $amp^R$ | This study |
| EJR298 | XL10-Gold pKSV7-$prfA^{G145S}$; $amp^R$ | This study |
| EJR299 | XL10-Gold pKSV7-$prfA^{V124A}$; $amp^R$ | This study |
| EJR305 | XL10-Gold pKSV7-$prfA^{wt}$; $amp^R$ | This study |
| EJR320 | XL10-Gold pKSV7-$prfA^{C229R}$; $amp^R$ | This study |
| EJR321 | XL10-Gold pKSV7-$prfA^{Y11D}$; $amp^R$ | This study |
| EJR322 | XL10-Gold pKSV7-$prfA^{V153M}$; $amp^R$ | This study |
| EJR331 | S17-1 pPL3e-$P_{actA}$-*lacZ*; $cat^R$ | This study |
| EJR334 | XL10-Gold pKSV7-$prfA^{D222E}$; $amp^R$ | This study |
| EJR335 | XL10-Gold pKSV7-$prfA^{V87I}$; $amp^R$ | This study |
| EJR336 | XL10-Gold pKSV7-$prfA^{D167Y}$; $amp^R$ | This study |
| EJR337 | XL10-Gold pKSV7-$prfA^{K64E}$; $amp^R$ | This study |
| EJR339 | XL10-Gold pKSV7-$prfA^{D33A}$; $amp^R$ | This study |
| EJR376 | XL10-Gold pKSV7-$prfA^{V96F}$; $amp^R$ | This study |
| EJR377 | XL10-Gold pKSV7-$prfA^{Y126F}$; $amp^R$ | This study |

| | | |
|---|---|---|
| EJR438 | DH5α pKSV7-*prfA*$^{C38A}$; amp$^{R}$ | This study |
| EJR439 | DH5α pKSV7-*prfA*$^{C38E}$; amp$^{R}$ | This study |
| EJR440 | DH5α pKSV7-*prfA*$^{C38S}$; amp$^{R}$ | This study |
| ***Listeria monocytogenes* strains** | | |
| ANG873 | EGD-e | [5] |
| BUG2214 | EGD-e Δ*prfA* | [6] |
| LJR105 | EGD-e *prfA*$^{D222E}$ Δ*gshF* pPL3e-$P_{actA}$-*lacZ*; ery$^{R}$ | This study |
| LJR106 | EGD-e *prfA*$^{V87I}$ Δ*gshF* pPL3e-$P_{actA}$-*lacZ*; ery$^{R}$ | This study |
| LJR108 | EGD-e *prfA*$^{D167Y}$ Δ*gshF* pPL3e-$P_{actA}$-*lacZ*; ery$^{R}$ | This study |
| LJR109 | EGD-e *prfA*$^{D33A}$ Δ*gshF* pPL3e-$P_{actA}$-*lacZ*; ery$^{R}$ | This study |
| LJR110 | EGD-e *prfA*$^{K64E}$ Δ*gshF* pPL3e-$P_{actA}$-*lacZ*; ery$^{R}$ | This study |
| LJR332 | EGD-e pIMK2-$P_{actA}$-*cat*; kan$^{R}$; short: $P_{actA}$-*cat* | This study |
| LJR333 | EGD-e pIMK2-$P_{hly}$-*cat*; kan$^{R}$ | This study |
| LJR334 | EGD-e Δ*prfA* pPL3e-$P_{plcA}$-$P_{prfA}$-*prfA*; ery$^{R}$; short: Δ*prfA* compl. or compl. | [4] |
| LJR345 | EGD-e pIMK2-$P_{hly}$-*cat* Suppressor *prfA*$^{L148F}$; kan$^{R}$ cat$^{R}$ | This study |
| LJR346 | EGD-e pIMK2-$P_{hly}$-*cat* Suppressor *prfA*$^{G145S}$; kan$^{R}$ cat$^{R}$ | This study |
| LJR350 | EGD-e Δ*gshF* | This study |
| LJR352 | EGD-e pPL3e-$P_{actA}$-*lacZ*; ery$^{R}$ | [4] |
| LJR354 | EGD-e Δ*gshF* pPL3e-$P_{actA}$-*lacZ*; ery$^{R}$ | This study |
| LJR362 | EGD-e pIMK2-$P_{actA}$-*cat* Suppressor *prfA*$^{V124A}$; kan$^{R}$ cat$^{R}$ | This study |
| LJR364 | EGD-e pIMK2-$P_{actA}$-*cat* Suppressor *prfA*$^{D222E}$; kan$^{R}$ cat$^{R}$ | This study |

| | | |
|---|---|---|
| LJR366 | EGD-e pIMK2-$P_{actA}$-*cat* Suppressor *rpoB*$^{S1128F}$; kan$^R$ cat$^R$ | This study |
| LJR367 | EGD-e pIMK2-$P_{actA}$-*cat* Suppressor *rpoC*$^{R328H}$; kan$^R$ cat$^R$ | This study |
| LJR368 | EGD-e pIMK2-$P_{hly}$-*cat* Suppressor *rpoB*$^{A1086E}$; kan$^R$ cat$^R$ | This study |
| LJR381 | EGD-e Suppressor *prfA*$^{F29L}$; kan$^R$ cat$^R$ | This study |
| LJR383 | EGD-e Suppressor *prfA*$^{Q35K}$; kan$^R$ cat$^R$ | This study |
| LJR389 | EGD-e *prfA*$^{wt}$ | This study |
| LJR391 | EGD-e *prfA*$^{V124A}$ | This study |
| LJR392 | EGD-e *prfA*$^{G145S}$ | This study |
| LJR393 | EGD-e *prfA*$^{L148F}$ | This study |
| LJR406 | EGD-e pIMK2-$P_{actA}$-*cat* Suppressor *prfA*$^{V96F}$; kan$^R$ cat$^R$ | This study |
| LJR408 | EGD-e pIMK2-$P_{actA}$-*cat* Suppressor *prfA*$^{V87I}$; kan$^R$ cat$^R$ | This study |
| LJR410 | EGD-e pIMK2-$P_{actA}$-*cat* Suppressor *prfA*$^{D167Y}$; kan$^R$ cat$^R$ | This study |
| LJR411 | EGD-e pIMK2-$P_{actA}$-*cat* Suppressor *prfA*$^{C229R}$; kan$^R$ cat$^R$ | This study |
| LJR422 | EGD-e pIMK2-$P_{hly}$-*cat* Suppressor *prfA*$^{D33A}$; kan$^R$ cat$^R$ | This study |
| LJR425 | EGD-e pIMK2-$P_{hly}$-*cat* Suppressor *prfA*$^{K64E}$; kan$^R$ cat$^R$ | This study |
| LJR441 | EGD-e *prfA*$^{D222E}$ pPL3e-$P_{actA}$-*lacZ*; ery$^R$ | This study |
| LJR442 | EGD-e *prfA*$^{V87I}$ pPL3e-$P_{actA}$-*lacZ*; ery$^R$ | This study |
| LJR444 | EGD-e *prfA*$^{D167Y}$ pPL3e-$P_{actA}$-*lacZ*; ery$^R$ | This study |
| LJR445 | EGD-e *prfA*$^{D33A}$ pPL3e-$P_{actA}$-*lacZ*; ery$^R$ | This study |

| | | |
|---|---|---|
| LJR446 | EGD-e *prfA*$^{K64E}$ pPL3e-$P_{actA}$-*lacZ*; ery$^R$ | This study |
| LJR447 | EGD-e *prfA*$^{F29L}$ pPL3e-$P_{actA}$-*lacZ*; ery$^R$ | This study |
| LJR448 | EGD-e *prfA*$^{Q35K}$ pPL3e-$P_{actA}$-*lacZ*; ery$^R$ | This study |
| LJR449 | EGD-e *prfA*$^{wt}$ pPL3e-$P_{actA}$-*lacZ*; ery$^R$ | This study |
| LJR450 | EGD-e *prfA*$^{V124A}$ pPL3e-$P_{actA}$-*lacZ*; ery$^R$ | This study |
| LJR451 | EGD-e *prfA*$^{G145S}$ pPL3e-$P_{actA}$-*lacZ*; ery$^R$ | This study |
| LJR452 | EGD-e *prfA*$^{L148F}$ pPL3e-$P_{actA}$-*lacZ*; ery$^R$ | This study |
| LJR463 | EGD-e *prfA*$^{F29L}$ Δ*gshF* | This study |
| LJR464 | EGD-e *prfA*$^{Q35K}$ Δ*gshF* | This study |
| LJR465 | EGD-e *prfA*$^{wt}$ Δ*gshF* | This study |
| LJR466 | EGD-e *prfA*$^{V124A}$ Δ*gshF* | This study |
| LJR467 | EGD-e *prfA*$^{G145S}$ Δ*gshF* | This study |
| LJR468 | EGD-e *prfA*$^{L148F}$ Δ*gshF* | This study |
| LJR489 | EGD-e pIMK2-$P_{hly}$-*cat* Suppressor *prfA*$^{Y126F}$; kan$^R$ cat$^R$ | This study |
| LJR499 | EGD-e *prfA*$^{F29L}$ Δ*gshF* pPL3e-$P_{actA}$-*lacZ*; ery$^R$ | This study |
| LJR500 | EGD-e *prfA*$^{Q35K}$ Δ*gshF* pPL3e-$P_{actA}$-*lacZ*; ery$^R$ | This study |
| LJR501 | EGD-e *prfA*$^{wt}$ Δ*gshF* pPL3e-$P_{actA}$-*lacZ*; ery$^R$ | This study |
| LJR502 | EGD-e *prfA*$^{V124A}$ Δ*gshF* pPL3e-$P_{actA}$-*lacZ*; ery$^R$ | This study |
| LJR503 | EGD-e *prfA*$^{G145S}$ Δ*gshF* pPL3e-$P_{actA}$-*lacZ*; ery$^R$ | This study |
| LJR504 | EGD-e *prfA*$^{L148F}$ Δ*gshF* pPL3e-$P_{actA}$-*lacZ*; ery$^R$ | This study |
| LJR505 | EGD-e *prfA*$^{D222E}$ | This study |
| LJR506 | EGD-e *prfA*$^{V87I}$ | This study |
| LJR508 | EGD-e *prfA*$^{D167Y}$ | This study |
| LJR509 | EGD-e *prfA*$^{D33A}$ | This study |
| LJR531 | EGD-e *prfA*$^{K64E}$ | This study |

| | | |
|---|---|---|
| LJR598 | EGD-e pIMK2-*P*$_{hly}$-*cat* Suppressor *rpoC*$^{V1060G}$; kan$^{R}$ cat$^{R}$ | This study |
| LJR603 | EGD-e pIMK2-*P*$_{actA}$-*cat* Suppressor *rpoC*$^{H1066P}$; kan$^{R}$ cat$^{R}$ | This study |
| LJR605 | EGD-e pIMK2-*P*$_{actA}$-*cat* Suppressor *prfA*$^{Y11D}$; kan$^{R}$ cat$^{R}$ | This study |
| LJR606 | EGD-e pIMK2-*P*$_{actA}$-*cat* Suppressor *prfA*$^{V153M}$; kan$^{R}$ cat$^{R}$ | This study |
| LJR623 | EGD-e *prfA*$^{D222E}$ ∆*gshF* | This study |
| LJR624 | EGD-e *prfA*$^{V87I}$ ∆*gshF* | This study |
| LJR626 | EGD-e *prfA*$^{D167Y}$ ∆*gshF* | This study |
| LJR627 | EGD-e *prfA*$^{D33A}$ ∆*gshF* | This study |
| LJR628 | EGD-e *prfA*$^{K64E}$ ∆*gshF* | This study |
| LJR629 | EGD-e *prfA*$^{C229R}$ | This study |
| LJR630 | EGD-e *prfA*$^{Y11D}$ | This study |
| LJR631 | EGD-e *prfA*$^{V153M}$ | This study |
| LJR632 | EGD-e *prfA*$^{V96F}$ | This study |
| LJR633 | EGD-e *prfA*$^{Y126F}$ | This study |
| LJR634 | EGD-e *prfA*$^{C229R}$ pPL3e-*P*$_{actA}$-*lacZ*; ery$^{R}$ | This study |
| LJR635 | EGD-e *prfA*$^{Y11D}$ pPL3e-*P*$_{actA}$-*lacZ*; ery$^{R}$ | This study |
| LJR636 | EGD-e *prfA*$^{V153M}$ pPL3e-*P*$_{actA}$-*lacZ*; ery$^{R}$ | This study |
| LJR637 | EGD-e *prfA*$^{V96F}$ pPL3e-*P*$_{actA}$-*lacZ*; ery$^{R}$ | This study |
| LJR638 | EGD-e *prfA*$^{Y126F}$ pPL3e-*P*$_{actA}$-*lacZ*; ery$^{R}$ | This study |
| LJR651 | EGD-e *prfA*$^{C229R}$ ∆*gshF* | This study |
| LJR652 | EGD-e *prfA*$^{Y11D}$ ∆*gshF* | This study |
| LJR653 | EGD-e *prfA*$^{V153M}$ ∆*gshF* | This study |
| LJR654 | EGD-e *prfA*$^{V96F}$ ∆*gshF* | This study |

| | | |
|---|---|---|
| LJR655 | EGD-e *prfA*$^{Y126F}$ Δ*gshF* | This study |
| LJR657 | EGD-e *prfA*$^{C229R}$ Δ*gshF* pPL3e-$P_{actA}$-*lacZ*; ery$^{R}$ | This study |
| LJR658 | EGD-e *prfA*$^{Y11D}$ Δ*gshF* pPL3e-$P_{actA}$-*lacZ*; ery$^{R}$ | This study |
| LJR659 | EGD-e *prfA*$^{V153M}$ Δ*gshF* pPL3e-$P_{actA}$-*lacZ*; ery$^{R}$ | This study |
| LJR660 | EGD-e *prfA*$^{V96F}$ Δ*gshF* pPL3e-$P_{actA}$-*lacZ*; ery$^{R}$ | This study |
| LJR661 | EGD-e *prfA*$^{Y126F}$ Δ*gshF* pPL3e-$P_{actA}$-*lacZ*; ery$^{R}$ | This study |
| LJR788 | EGD-e *prfA*$^{C38A}$ | This study |
| LJR789 | EGD-e *prfA*$^{C38E}$ | This study |
| LJR790 | EGD-e *prfA*$^{C38S}$ | This study |
| LJR818 | EGD-e *prfA*$^{C38A}$ pPL3e-$P_{actA}$-*lacZ*; ery$^{R}$ | This study |
| LJR819 | EGD-e *prfA*$^{C38E}$ pPL3e-$P_{actA}$-*lacZ*; ery$^{R}$ | This study |
| LJR820 | EGD-e *prfA*$^{C38S}$ pPL3e-$P_{actA}$-*lacZ*; ery$^{R}$ | This study |
| LJR850 | EGD-e *prfA*$^{C38A}$ Δ*gshF* | This study |
| LJR851 | EGD-e *prfA*$^{C38E}$ Δ*gshF* | This study |
| LJR852 | EGD-e *prfA*$^{C38S}$ Δ*gshF* | This study |
| LJR853 | EGD-e *prfA*$^{C38A}$ Δ*gshF* pPL3e-$P_{actA}$-*lacZ*; ery$^{R}$ | This study |
| LJR854 | EGD-e *prfA*$^{C38E}$ Δ*gshF* pPL3e-$P_{actA}$-*lacZ*; ery$^{R}$ | This study |
| LJR855 | EGD-e *prfA*$^{C38S}$ Δ*gshF* pPL3e-$P_{actA}$-*lacZ*; ery$^{R}$ | This study |

**Table S2: Primers used in this study**

| **Number** | **Name** | **Sequence** |
|---|---|---|
| JP3 | $P_{actA}$ fw for *lacZ* | AAAGGATCCGTTTACCGATGCGAAAAAAGCG |
| JP4 | $P_{actA}$ rev for *lacZ* | TTTGTCGACCACTTATACTCCCTCCTCGT |
| JR257 | *prfA* rev | ACGCGTCGACCGAATAAAATATAAAACAGTATTCCTC |
| JR258 | *prfA* up fw | ACGCGTCGACGGAATAAGCCAATAAAGAACTCTG |
| JR261 | *prfA* down rev | CGGGGTACCGTGGAGGCTGATTATGTCAAAC |
| JR295 | $P_{actA}$ fw for *cat* | CCGGAATTCGCAGCGAAAGATTTATATGGTG |

| | | |
|---|---|---|
| JR296 | $P_{actA}$ rev for *cat* | ATTAAAGTTCATTTATACTCCCTCCTCGTGATAC |
| JR297 | *cat actA* fw | GAGGGAGTATAAATGAACTTTAATAAAATTGATTTAG ACAATT |
| JR298 | *cat* rev | CCGGAATTCTTATAAAAGCCAGTCATTAGGCC |
| JR299 | $P_{hly}$ fw for *cat* | CCGGAATTCGTGACTTTTATGTTGAGGCATTAA |
| JR300 | $P_{hly}$ rev for *cat* | ATTAAAGTTCATGGGTTTCACTCTCCTTCTAC |
| JR301 | *cat hly* fw | AGAGTGAAACCCATGAACTTTAATAAAATTGATTTAG ACAATT |
| JR312 | $P_{plcA}$ fw | CGGGGTACCGAAATTCGCTTCTAAAGATGAAACG |
| JR313 | $P_{plcA}$ rev | GTTTTTTATGATGTTTAATGTGCGTTGCAAATAATTCT TATA |
| JR314 | *prfA* fw | GCAACGCACATTAAACATCATAAAAAACAATAAATTC AACTAAC |
| JR319 | *gshF* up fw | AAAGGTACCCACCGTGAAAATGTCTTCTATC |
| JR320 | *gshF* up rev | TAAAATCTTATCATCAAGCATGGTCATATCAAGTTTT A |
| JR321 | *gshF* down fw | ACCATGCTTGATGATAAGATTTTAGATTTCTTATTTGA CTAAA |
| JR322 | *gshF* down rev | TTTGTCGACCGCTTGGGATTCGTCATGC |

**Table S3: Frequency of identified PrfA mutations**

| Mutation | Frequency | Mutation | Frequency |
|---|---|---|---|
| Y11D | 1 | Y126F | 1 |
| F29L | 2 | L140F | 3 |
| D33A | 2 | G145S | 7 |
| Q35K | 1 | G145C | 1 |
| Q35R | 1 | G145V | 2 |
| I45N | 1 | G145A | 2 |
| I45S | 2 | L148F | 1 |
| I45T | 4 | V153M | 1 |
| K64E | 1 | G155S | 5 |

| | | | |
|---|---|---|---|
| V87I | 1 | D167Y | 2 |
| A94T | 7 | D167N | 1 |
| V96F | 3 | P219S | 2 |
| V96L | 1 | D222E | 1 |
| V124A | 2 | C229R | 1 |

**Table S4: Overview of PrfA* phenotypes**

| Mutation | PlcB 30°C Cello | Growth LSM Glc | Growth LSM Glc1P | Glutathione-dependence | Sugar-repression $P_{actA}$? | Intracellular replication |
|---|---|---|---|---|---|---|
| Y11D | - | wt | + | +++ | - | n.d. |
| F29L | ++ | wt | +++ | + | + | n.d. |
| D33A | + | wt | + | +++ | ++ | n.d. |
| Q35K | ++ | wt | ++++ | ++ | - | n.d. |
| K64E | ++ | wt | ++ | ++ | - | n.d. |
| V87I | - | wt | ++ | ++++ | - | - |
| V96F | +++ | wt | +++ | +++ | -/+ | - |
| V124A | - | wt | ++ | +++ | - | - |
| Y126F | +++ | wt | +++ | - | + | n.d. |
| G145S | +++ | - | +++ | - | - | +++ |
| L148F | +++ | - | +++ | +++ | +++ | - |
| V153M | - | wt | + | - | - | n.d. |
| D167Y | -/+ | wt | - | ++++ | +++ | - |

| | | | | | | |
|---|---|---|---|---|---|---|
| D222E | ++ | wt | ++ | +++ | ++ | - |
| C229R | -/+ | wt | + | ++++ | + | n.d. |

wt – wildtype-like, Glc – Glucose; Glc1P – Glucose-1-phosphate, n.d. – not determined

### Supplemental Figures

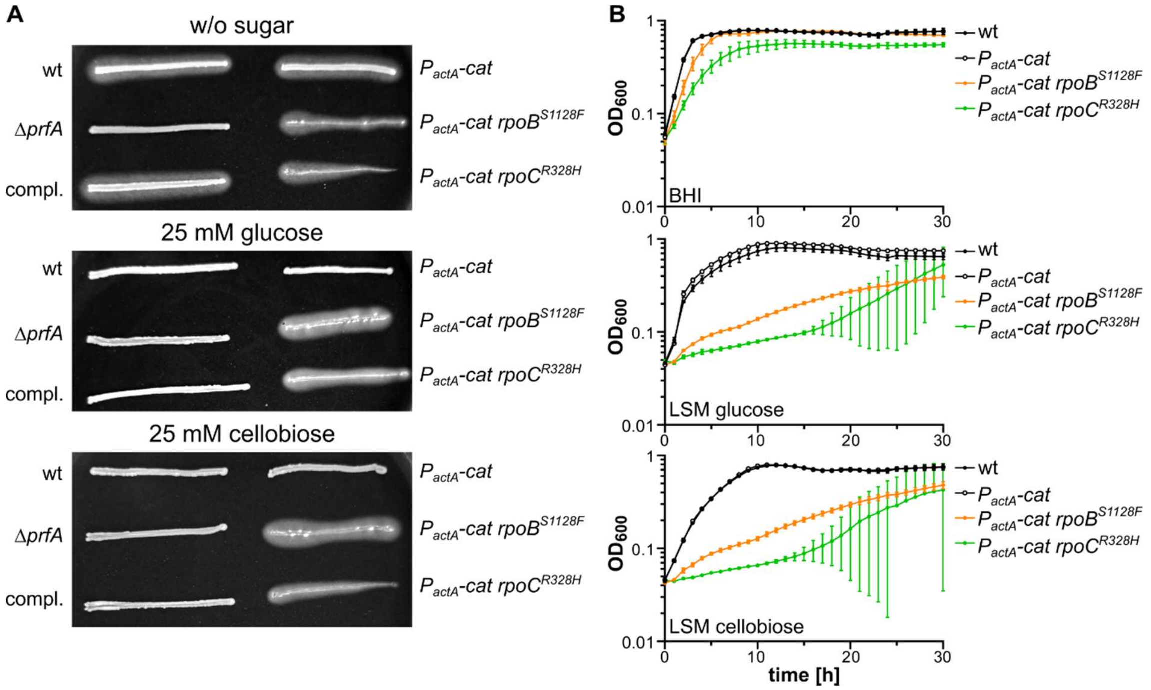


**Fig. S1: Impact of RpoB and RpoC mutation on PlcB activity and growth.** (A) PlcB assay. *L. monocytogenes* strains EGD-e (wt), LJR332 ($P_{actA}$-*cat*), LJR366 ($P_{actA}$-*cat rpoB*$^{S1128F}$) and LJR367 ($P_{actA}$-*cat rpoC*$^{R328H}$) were streaked onto LB agar containing activated charcoal, egg yolk, and where indicated, 25 mM glucose or cellobiose. The *prfA* deletion strain and the *prfA* complementation strain (compl.) were used as reference. The plates were incubated for a day at 37°C and representative images of at least three independent experiments are shown. (B) Overnight cultures of the indicated *L. monocytogenes* strains were diluted in fresh BHI and grown until an $OD_{600}$ of 0.3. 2 ml of the cultures were harvested, washed twice in BHI, LSM glucose or LSM cellobiose and resuspended in 1 ml of the corresponding medium. The cell suspension was adjusted to an $OD_{600}$ of 0.1 and transferred into wells of a 96-well plate. Growth of the strains was measured every 15 min for 30 h. Averages and standard deviations of three independent experiments are plotted.

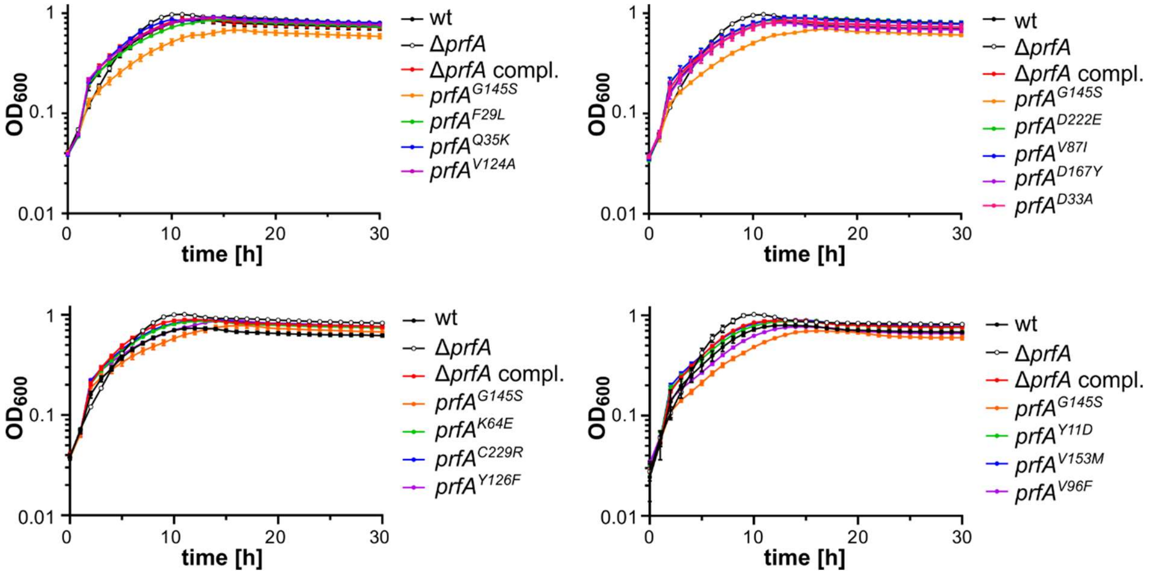


**Fig. S2: Effect of PrfA mutations on growth in LSM glucose.** Overnight cultures of the indicated *L. monocytogenes* strains were diluted in fresh BHI and grown until an $OD_{600}$ of 0.3. 2 ml of the cultures were harvested, washed twice in LSM glucose and resuspended in 1 ml LSM glucose. The cell suspension was adjusted to an $OD_{600}$ of 0.1 and transferred into wells of a 96-well plate. Growth of the strains was measured every 15 min for 30 h. Averages and standard deviations of three independent experiments are plotted.

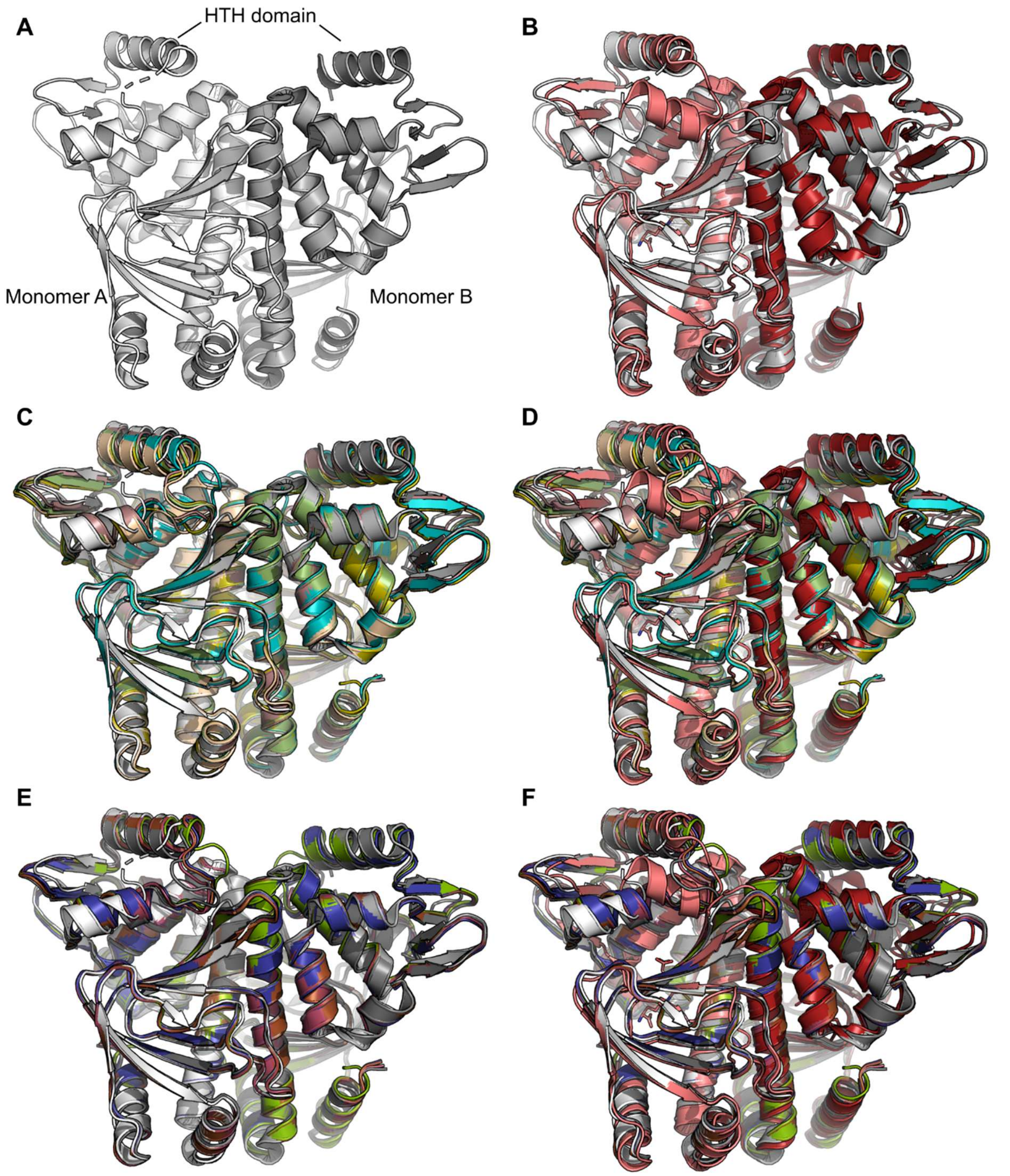


**Fig. S3: AlphaFold structure predictions of Glutathione-independent PrfA* variants.** (A) Crystal structure of apo-PrfA homodimer (PDB: 1OMI; grey). Individual monomers are colored in light and dark grey. (B) Crystal structures of apo-PrfA homodimer and glutathione-bound PrfA (PDB: 5LRR; red) [7]. The individual monomers of the glutathione-bound PrfA are colored in light and dark red. (C) The top 5 highest-scoring predictions for the homodimer of PrfA$^{Y126F}$ were superimposed on the crystal structure of apo-PrfA. (D) The top 5 highest-scoring predictions for the homodimer of PrfA$^{Y126F}$ were superimposed on the crystal structure of apo-PrfA and glutathione-bound PrfA. (E) The top 5 highest-scoring predictions for the homodimer of PrfA$^{V153M}$ were superimposed on the crystal structure of apo-PrfA. (F) The top 5 highest-scoring predictions for the homodimer of PrfA$^{V153M}$ were superimposed on the crystal structure of apo-PrfA and glutathione-bound PrfA.
